# JACUZI-SD: An automated, high-throughput, minimally stressful approach to sleep depriving larval zebrafish

**DOI:** 10.1101/2025.04.03.647099

**Authors:** Leah J. Elias, Harrison Khoo, François Kroll, Caroline Zhang, Soojung C. Hur, Jason Rihel, Seth Blackshaw

## Abstract

While sleep deprivation broadly disrupts health and well-being, the neural and molecular mechanisms that signal increased sleep pressure remain poorly understood. A key obstacle to progress is the fact that traditional methods for inducing sleep deprivation (SD) in animal models often introduce confounding stress or disrupt circadian rhythms. Here, we present JACUZI-SD (Jetting Automated Currents Under Zebrafish to Induce Sleep Deprivation), a fully automated, high-throughput system designed to induce sleep deprivation in larval zebrafish with minimal stress. By delivering randomized water pulses via a custom milli-fluidic device integrated with a 96-well plate and controlled by an Arduino, JACUZI-SD promotes wakefulness during the natural dark cycle without the stress associated with existing SD methods. Our results demonstrate that JACUZI-SD reduces total sleep time by 41-64% and elicits a robust rebound sleep characterized by increased sleep bout length following deprivation. Importantly, this method avoids activating the hypothalamic-pituitary-interrenal (HPI) stress axis, as evidenced by reduced stress marker expression compared to other deprivation methods. Additionally, the system reliably activates established sleep pressure pathways, including the upregulation of *galanin* in the neurosecretory preoptic area, while also revealing biologically relevant inter-individual variability in homeostatic rebound responses. JACUZI-SD provides a powerful, minimally invasive platform for dissecting the neural and molecular underpinnings of sleep homeostasis in vertebrates.

## Introduction

The conservation of sleep across animal species is remarkable considering the opportunity cost of forgoing evolutionarily beneficial behaviors (feeding, reproduction) while making oneself vulnerable to predators. That sleep is critical for life is further demonstrated by the many detrimental effects of its absence^1^. Indeed, inadequate sleep is one of the greatest burdens on our healthcare system, impacting an estimated 33.2% of Americans^2^. However, the biological manifestation of this fundamental homeostatic need remains unknown. The homeostatic drive to sleep is called sleep pressure, which increases with wakefulness and dissipates with sleep independently of the circadian drive for arousal^2–4^. Isolating the biological underpinnings of sleep pressure from other variables is challenging, as experimental sleep deprivation (SD) inherently induces the confound of changing the waking experience^1, 5^. Stress in particular is a common confound: while SD is inherently stressful, approaches to sleep depriving animals are often excessively so, promoting wakefulness by mimicking an ethological threat such as a predator or impending physical injury^5^. Furthermore, efforts to minimize stress are often labor-intensive and low-throughput, whereas high-throughput automated approaches are likely to be more stressful because they cannot be tuned to the minimum stimulation each animal requires to be kept awake.

In recent years, larval zebrafish have demonstrated great potential as model organism for sleep research^6–14^. In addition to their amenability to *in vivo* imaging and transgenic manipulation, larval zebrafish exhibit diurnal sleep-wake behavior that is governed by neuronal machinery much conserved with that of mammals^6^. A hypothalamic regulatory center controls the sleep-wake brain state using many of the same neuropeptides and transmitters critical for mammalian sleep function^6, 13^. Remarkably, these cells and molecules operate within an experimentally tractable ∼100,000 neuron brain, making it feasible to identify small functional subtypes that would be obscured in the mammalian brain. Perhaps most powerfully, clutches of hundreds of siblings facilitate high-throughput studies, which are critical to capture the biological variability in these homeostatic behaviors.

However, SD methods in larval zebrafish with sufficient efficacy and throughput have been challenging to establish without introducing confounds. Mechanical methods, involving shaking or rotating, while high-throughput, preclude simultaneous behavioral tracking and can cause stress or even injury^8, 15^. It is therefore important to distinguish whether the behavioral rebound response is stress-driven or sleep pressure-driven by conducting the manipulation during the light cycle when sleep would not be disturbed.^16^ While this approach rules out the behavioral rebound being driven by stress, few studies directly assess the neurobiological stress pathways themselves. Applying or simulating (optical treadmill ^13^) water flow^17^ in groups of fish is likely less stressful, but individual behavioral data is still lost. Extending or altering the light cycle, while minimally invasive and efficacious to promote wakefulness^12, 18^ ^19^, does manipulate the circadian clock. This makes it impossible to isolate the neurobiological impact of heightened homeostatic sleep pressure (“Process S”) from the circadian drive for wakefulness (“Process C”) ^3^. Lastly, many of these automated stimuli can ultimately be ignored in the face of extreme sleep pressure. For example, one study found that achieving behaviorally significant sleep deprivation with gentle rotation to promote continuous swimming requires three days of sleep deprivation^8^, while other comparable methods produce rebound after 4-6 hours of SD. On the other hand, forced movement methods like paintbrush-mediated gentle handling^11^ effectively maintain wakefulness but are frustratingly low-throughput and labor intensive for a high-throughput model system like larval zebrafish. Furthermore, the paintbrush resembles a looming, predator-like stimulus that is often used to experimentally induce stress in larvae^20^. In addition to introducing the potential confound of stress, paintbrush manipulation prevents video behavior tracking and can introduce experimenter-dependent variation. The confounding variables of stressors or zeitgebers in efficacious SD methods make it impossible to isolate the neurobiological changes associated specifically with homeostatic sleep pressure, as opposed to those driven by stress or circadian arousal.

There is therefore a critical need for a high-throughput, minimally stressful, automated approach to sleep deprive larval zebrafish. Here we present JACUZI-SD (Jetting Automated Currents Under Zebrafish to Induce Sleep Deprivation), a fully automated, minimally stressful system to sleep deprive up to 96 (or 48 with within-plate controls) larvae by randomized water pulses during simultaneous tracking of sleep-wake behavior in the widely used ZebraBox (ViewPoint LifeSciences). By using naturalistically non-threatening water flow to maintain wakefulness, JACUZI-SD isolates the accumulation of homeostatic sleep pressure from common confounds of experimental SD such as predator-like stressors, manipulation of the circadian clock, and inter-experimenter variability.

## Results

### JACUZI-SD enables high-throughput, automated sleep deprivation of larval zebrafish during simultaneous tracking of sleep-wake behavior

To uncover neural mechanisms of homeostatic sleep pressure, there is a critical need for novel approaches to sleep depriving animals in a high-throughput manner that minimizes confounding variables such as stress or manipulations to the circadian clock. We sought to develop such an approach in larval zebrafish, which are amenable to high-throughput behavioral studies due to their large clutch sizes and rapid development of diurnal sleep-wake behaviors and neuronal machinery. We decided to use water pulses as a wake-promoting stimulus because they are not inherently threatening in the natural environment and changes in water flow are sufficient to keep fish awake^17^. Furthermore, we wanted the sleep deprivation to be compatible with simultaneous video tracking of larval behavior, and water pulses offered an optically transparent stimulus that would minimize interference with behavioral acquisition. As larval zebrafish are prone to habituate to repeated stimulation^21^, we used an Arduino-compatible submersible water pump to facilitate randomization of water pulse frequency and duration with random number generators (**Figure 1A-C**). To combat increasing sleep pressure, the pulses increased in frequency and duration with each hour of SD by decreasing the range supplied to the random number generator that calculates inter-pulse interval and increasing the range supplied to the random number generator that calculates the duration the pump is powered on. The full Arduino code is provided in **Supplemental Document 1**.

**Figure 1:**
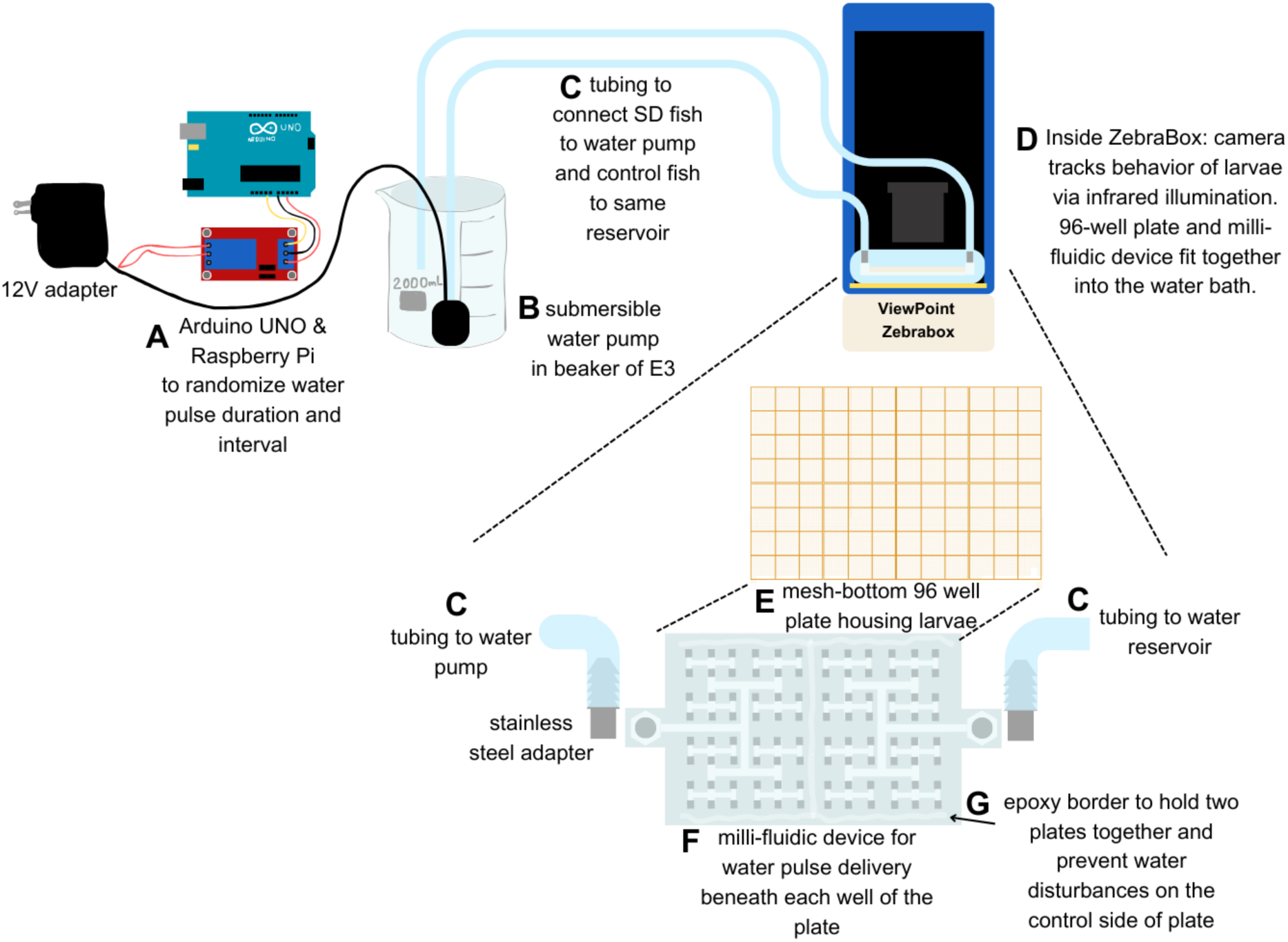
JACUZI-SD Diagram. **A)** Arduino UNO wired to Raspberry Pi. Pulse duration and inter-pulse interval are randomized to minimize habituation to the stimulus. Pulse duration increases and inter-pulse interval decreases with each hour of sleep deprivation to combat increasing sleep pressure and minimize habituation. **B)** Submersible water pump in a beaker of E3 attached to **C)** nontoxic plastic tubing, which is threaded through the back panel of the ZebraBox, and the gap generated is tightly covered with light-blocking fabric. **D)** Within the ZebraBox, infrared illumination allows high-resolution camera to record sleep behavior of 96 larvae individually housed in a mesh-bottom 96 well plate secured into the milli-fluidic device base. The water bath sits on top of an IR and LED light source to provide 14hr:10hr light:dark cycles. **E)** 3D-printed mesh bottom 96 well plate adapted from Kroll et al., 2025^22^. **F)** Stereolithography-printed milli-fluidic device secured beneath the 96 well plate provides channels to deliver water pulses beneath each larva. The entire arena is symmetrical to prevent any baseline behavioral differences across the plate. Stainless steel adaptor screws into the milli-fluidic device and tubing is attached to barbs. Parafilm is used around this junction to minimize loss of water flow.

To deliver water pulses to larvae housed in individual wells of a 96-well plate typically used for behavior tracking, we developed a milli-fluidic device geometrically compatible with a mesh-bottomed 96 well plate^22^ and ZebraBox (**Figure 1C-F; Figure S1**). The milli-fluidic device receives water from a shared reservoir through two mirrored inlets, which symmetrically distribute flow across a channel network. One inlet delivers water from the left middle channel to 48 wells, while the other does so from the right middle channel to the remaining 48 wells, directing pulses to the mesh bottom of each of the 96 wells (**Figure 1E,F; Figure S1**). The lefthand side of the plate is connected to the submersible water pump while the righthand side of the plate is connected to the same external reservoir of embryo water without a water pump. This setup harmonizes the environment between experimental groups as our rested control larvae are housed within the same ZebraBox, and backfilled from the same external reservoir, as SD larvae. To prevent the pulses on the lefthand side from creating a wave that would disrupt the sleep of rested controls, we added a water-tight epoxy barrier (**Figure 1G**) between groups, which also serves to affix the milli-fluidic device to the 96-well plate to form the “jacuzzi”. Further details on the JACUZI-SD device and experimental set up can be found in Methods.

### JACUZI-SD sleep deprives larvae

To test the function of JACUZI-SD, we measured sleep and activity behavior of larvae before, during, and after six hours of water pulse sleep deprivation. Larvae were raised on a 14hr:10hr reverse light-dark cycle and plated in the “jacuzzi” at 4 dpf. After 24 hours of habituation, baseline sleep behavior was collected on the second dark cycle. The third dark cycle consisted of 6 hours of JACUZI-SD water pulses followed by 4 hours of recovery. The sleep and activity traces of a single representative experiment are shown in **Figure 2A** and **2B**, respectively, while D-H are summary data of multiple experiments. To assess whether our design was sufficient to significantly disrupt sleep compared to rested controls, we compared sleep and activity during the pulses to the behavior during the equivalent 6-hour time window during the baseline night (**Figure 2C-H**). To control for individual variability, each animal is normalized to itself: its percent change is calculated from its own baseline. Because we observe a progressive reduction in sleep time in control larvae (visualized from 4dpf to 5dpf and 5dpf to 6dpf in **Figure 2A**), we adjust for these developmental sleep changes by subtracting out the average percent change of control animals from all animals to obtain normalized percent change from baseline day (**Figure 2A** yellow notations. See also Methods section “Collection, Normalization, and Statistical Analysis of Sleep-Wake Data”). Larvae exposed to 6h JACUZI-SD had a 64% average reduction in their total sleep time (sleep is here defined as one minute of inactivity) from baseline night compared to rested clutchmate controls (**Figure 2D**).

**Figure 2:**
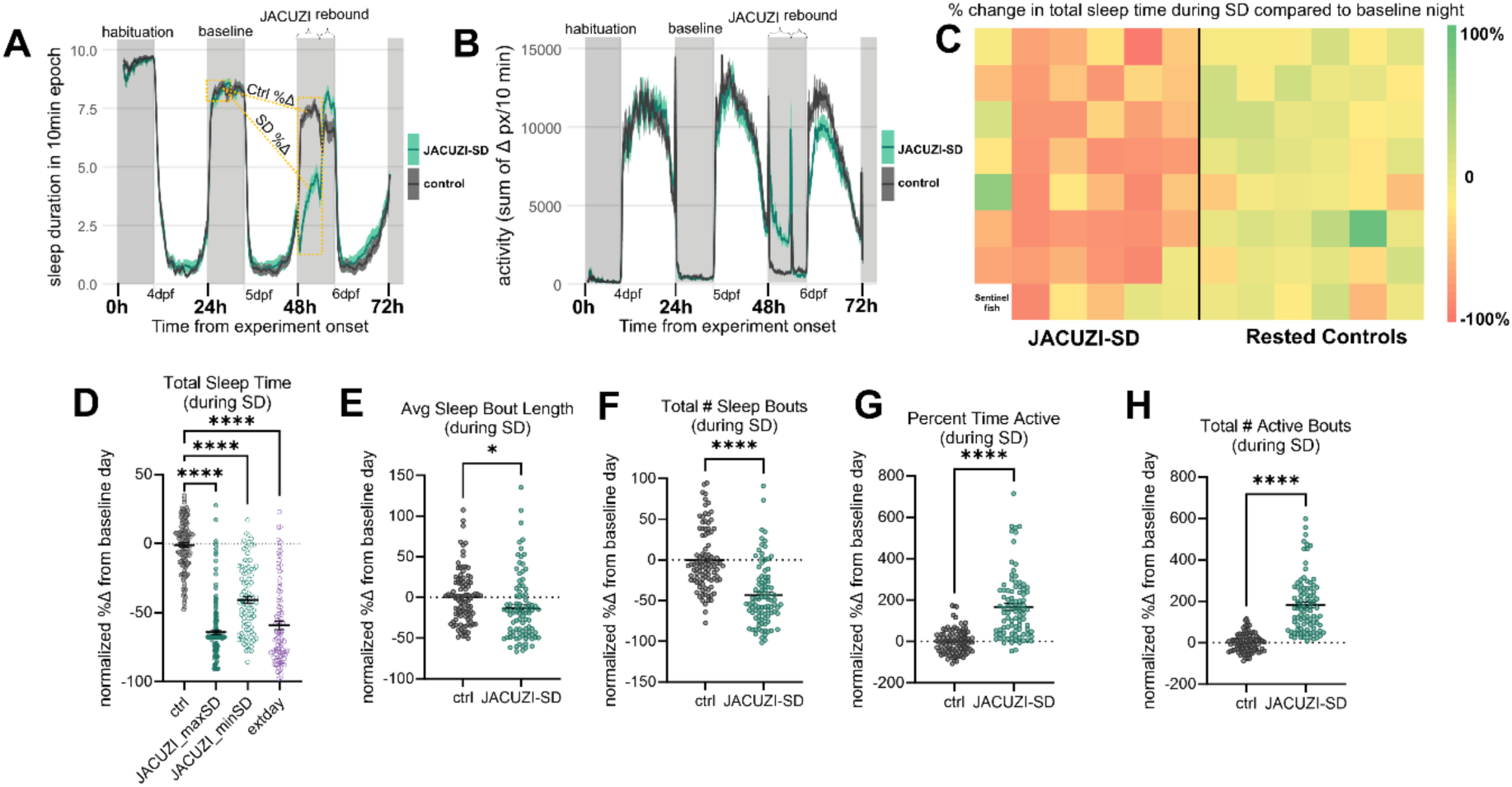
JACUZI-SD is an effective, high-throughput, automated approach to prevent sleep during the natural dark cycle. **A)** Average sleep trace of 95 larvae (*n* = 47 SD, 48 ctrls) during a single representative JACUZI-SD experiment. Sleep is calculated as 1 minute of inactivity and plotted here as average sleep duration in 10 minute epoch. Aqua trace is JACUZI-SD group, gray trace is control group. Dark cycle is represented by gray bars; light cycle represented by white bars. Normalization method is schematized by yellow notations: percent change for a given behavioral parameter is calculated for each animal from baseline night to the corresponding period on SD night, either 6h SD (shown here) or rebound period. The average percent change among controls is subtracted from each animal to obtain “normalized %Δ from baseline day” plotted in D-H. **B)** Average activity trace of 95 larvae (*n* = 47 SD, 48 ctrls) during a single representative JACUZI-SD experiment. Activity is calculated as the sum of pixel changes from one frame to the next in a 10 minute epoch. **C)** Heat map representing sleep deprivation of individual larvae across the 96 well plate for a representative JACUZI-SD experiment. Values calculated as percent change in total sleep time during sleep deprivation as compared to baseline night. Warmer colors are a decrease in sleep time, cooler colors represent an increase in sleep time. **D)** Total sleep time during SD in rested controls, approximated maximum SD (not removing pulse frames), approximated minimum SD (excluding water pulse frames), and larvae exposed to 4 hours of extended day (*p<*0.0001; *n* = 137, 140, 92, 94; One-Way ANOVA followed by Tukey’s Multiple Comparisons Test). **E)** Normalized percent change in sleep bout length during JACUZI-SD pulses (*p*=0.0141; *n* = 96, 92; Unpaired t-Test) **F)** Normalized percent change in number of sleep bouts during JACUZI-SD pulses (*p<*0.0001, *n* = 95, 92; Unpaired t-Test) **G)** Normalized percent change in total number of active bouts during JACUZI-SD pulses (*p<*0.0001, *n* = 95, 91; Unpaired t-Test). **H)** Normalized percent change in percentage time active during JACUZI-SD pulses (*p<*0.0001, *n* = 94, 91; Unpaired t-Test). Figures **E-H** represent two experimental replicates. *(*p*<0.05; *\*\*p*<0.01*; ***p*<0.001*; ****p<*0.0001)

As the water pulses generate movement of the larvae that is not always indicative of wakefulness, an observed 64% reduction in total sleep time represents the maximum estimated SD, as this calculation includes forced movements by the water pulse that fail to awaken the larva. We therefore sought a way to exclude from the analysis the video frames in which the pulses cause water movement and exclusively analyze volitional movements during the SD window (**Figure S2**). Since this analysis makes the conservative assumption that larvae are not awakened during the water movements (e.g., a sleeping fish being swirled without awakening) but also excludes true positive movement during the pulses (e.g. a fish exhibiting wakeful, volitional movement while the water flow is running), this analysis represents a minimum estimate of SD received by the larvae. With these calculations, we conclude that JACUZI-SD drives an average of 41-64% further reduction in total sleep time relative to baseline sleep, beyond what is observed in rested controls due to developmental changes (**Figure 2D**, further details in Methods). JACUZI-SD performs approximately as well as extended day at maintaining wakefulness without introducing a light stimulus that will alter the circadian clock phase, as pulses occur during the natural dark cycle (**Figure 2D**). For the remainder of behavioral analysis during the pulses, we apply the more conservative analysis and exclude the water pulse frames.

Figure 2C depicts the percent reduction in total sleep time during the pulses compared to baseline night across the plate as a heatmap. Even with this more conservative analysis, we see a 13±5% reduction in sleep bout length (Figure 2E) and 43±6% reduction in the number of sleep bouts beyond the developmental sleep changes observed in rested controls (Figure 2F). The overall activity of the larvae is also increased: they exhibit a 167±17% increase in the percentage of time active (Figure 2G) and 181±15% increase in number of active bouts (Figure 2H) during the pulses compared to their baseline night beyond what is observed in rested controls.

## JACUZI-SD induces rebound sleep

Since sleep pressure is homeostatically regulated, one indicator of successful SD is a consequent increase in sleep time or consolidation, termed “recovery” or “rebound” sleep. For this reason, we used the last 3 hours of the natural dark cycle (excluding a one-hour transition window from SD to sleep, as the experimenter enters the room to refill water levels immediately following SD) as a rebound period in which to analyze the changes in sleep architecture associated with having been exposed to 6 hours of JACUZI-SD water pulses. Larvae exposed to 6 hours of JACUZI-SD water pulses had an 11±3% percent increase in total sleep time (Figure 3A) and 29±6% increase in sleep bout length (Figure 3B) compared to rested controls. SD larvae also exhibited a 21±6% reduction in sleep bout number (Figure 3C), suggesting that the SD larvae exhibit a consolidated rebound sleep after JACUZI-SD, characterized by an increase in sleep due to lengthened sleep bouts relative to their baseline sleep that is not explained by developmental changes in sleep architecture seen in rested controls. In addition, SD larvae exhibit a 67±23% reduction in percentage time active (Figure 3D) and a 57±17% decrease in active bout number (Figure 3E) during the rebound period. After manipulation, a slight reduction in sleep is often observed in control animals (as in the experiment represented in Figure 2A sleep trace). We conclude this is due to experimenter intervention after pulses, in which the experimenter enters the room to check on the larvae and top off water levels, possibly promoting arousal. This manipulation is equal across SD and control animals, and therefore is accounted for in our method of normalization.

**Figure 3:**
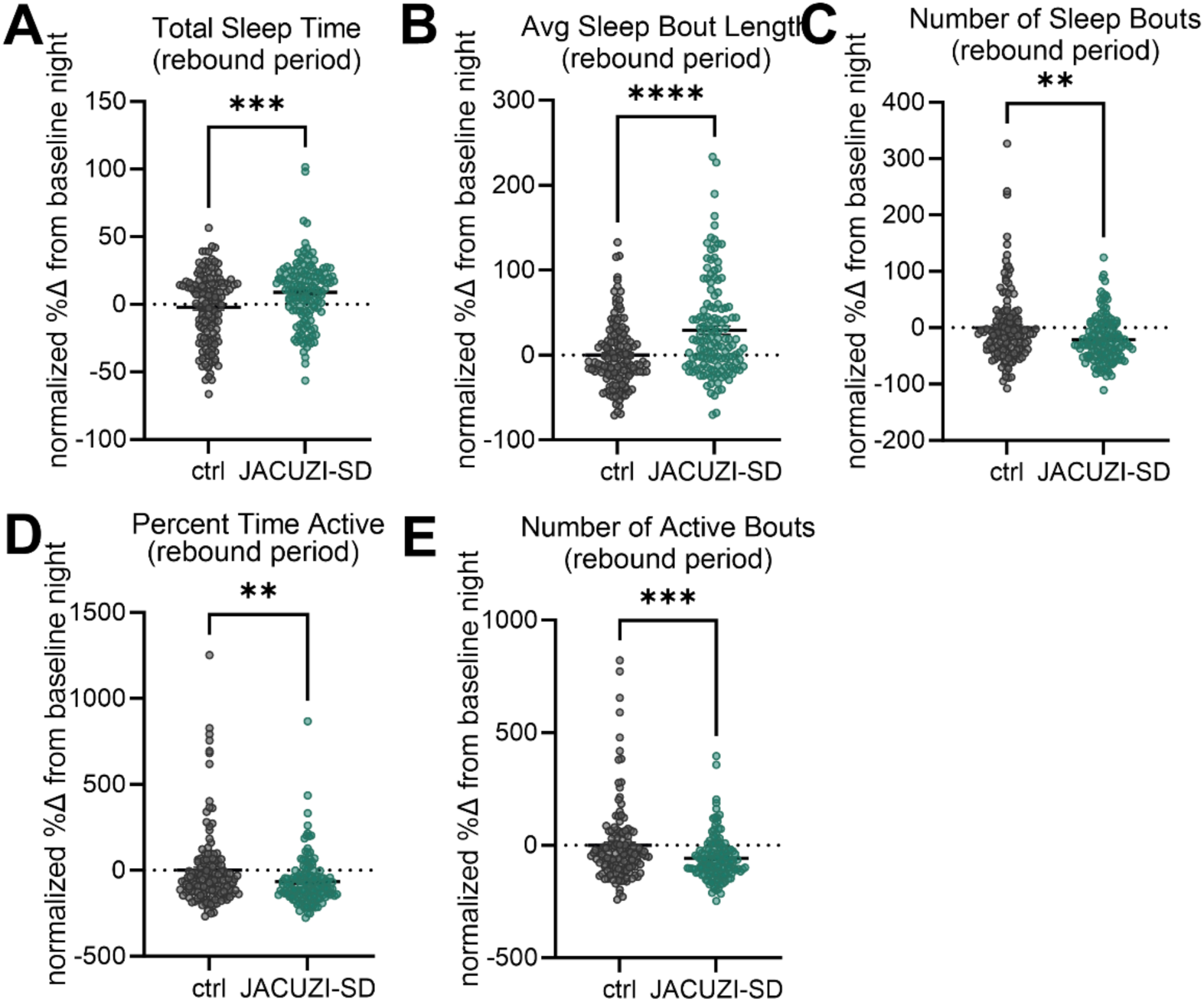
JACUZI-SD induces rebound sleep. Normalized percent change in A) total sleep time (*p=*0.0002; *n* = 140, 139), B) sleep bout length (*p=*0.0001; *n* = 140, 138), C) number of sleep bouts (*p=*0.0013; *n* = 141, 138), D) percent time active (*p=*0.0039; *n* = 140, 137), and E) number of active bouts (*p=*0.0009, *n* = 140, 137) during the 4h rebound period compared to the 4h baseline period 24h prior. *(*p<0.05; **p<0.01; ***p<0.001; ****p<0.0001;* Unpaired t-Tests)

To evaluate whether sleep loss or the stress of manipulation is the primary driver of this rebound response, we delivered 6 hours of JACUZI-SD pulses during the light cycle and measured rebound behavior. There was no significant change in percentage time active, number of active bouts, total sleep time, sleep bout length, or sleep bout number during the remainder of the light cycle after JACUZI-SD from baseline day compared to undisturbed clutchmate controls. (**Figure S3**)

### JACUZI-SD minimizes stress as a confounding variable

While these experiments establish that sleep rebound was not likely driven by the stress of the JACUZI pulse manipulation, we wanted to formally measure the neurobiological level of stress induced by the manipulation itself. To test our hypothesis that water pulses would not be stressful to the larvae, we examined the activation of the hypothalamic pituitary interrenal (HPI) stress axis when the JACUZI-SD pulses were applied during the daytime or nighttime (Figure 4A). Using HCR fluorescent *in situ* hybridization, we measured *fosab* induction in *corticotropin hormone (crh)+* neurons of the neurosecretory preoptic area as well as *pomc+* (which encodes adrenocorticotropic hormone (ACTH)) cells of the anterior pituitary (Figure 4B**,C**). To validate this approach to measuring stress, we included a positive control stressor known to induce cortisol release (“mesh stress”), in which larvae are intermittently withdrawn from water in a mesh cell strainer^23, 24^. Indeed, we observed strong HPI axis activation in these larvae (Figure 4D-G).

**Figure 4:**
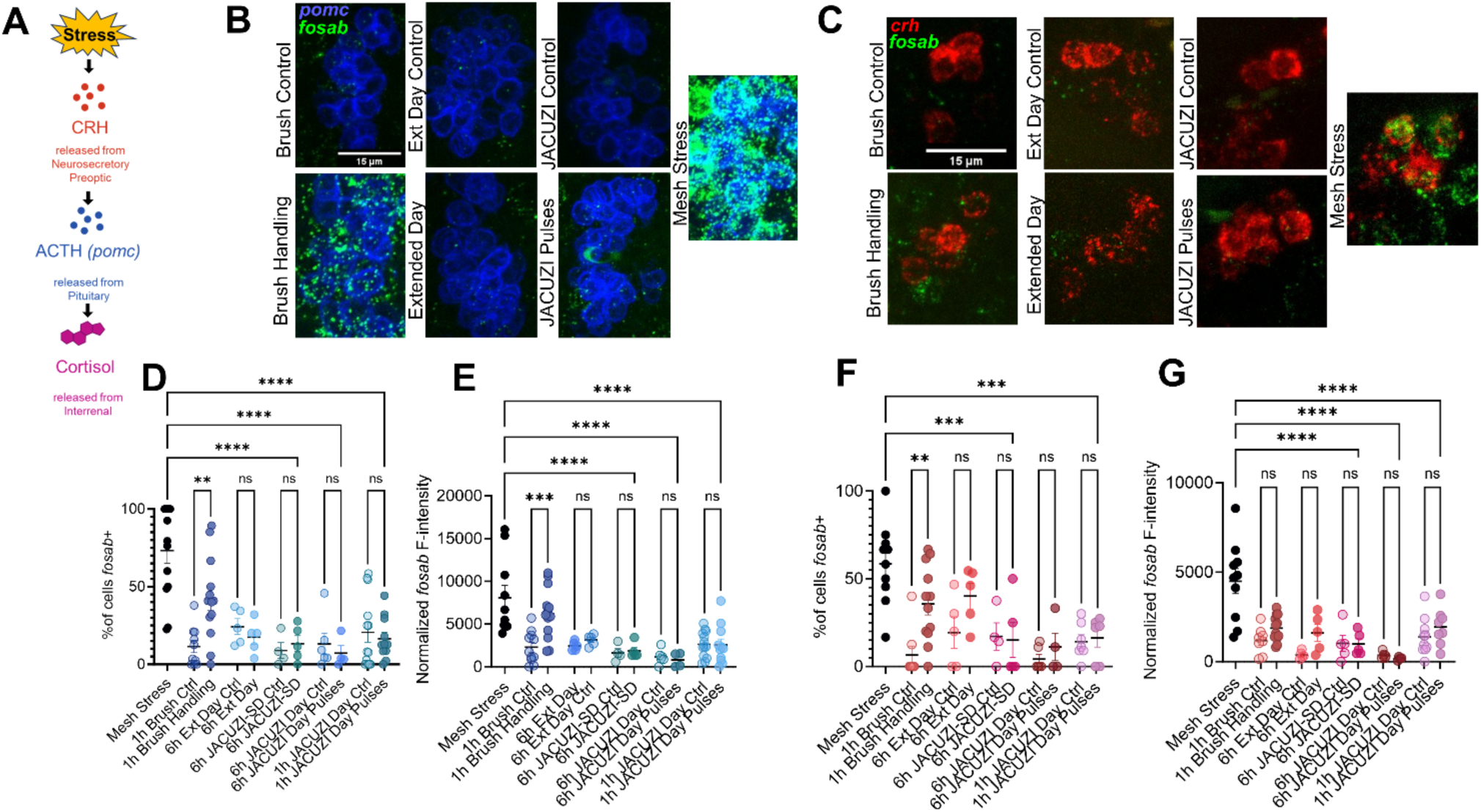
JACUZI-SD is less stressful than comparable methods of SD. **A)** Schematic depicting the zebrafish HPI axis, adapted from Eachus et al., 2022^19^. **B)** Representative images of double fluorescent *in situs* (HCR) for *acth* (blue) and *fosab* (green) in the anterior pituitary and **C)** *crh* (red) and *fosab* (green) in the neurosecretory preoptic of controls from paintbrush manipulation and clutchmates exposed to one hour of paintbrush gentle handling; controls from extended day SD and clutchmates exposed to 6h extended day SD; controls from 1h JACUZI in the ZebraBox and clutchmates exposed to 1h JACUZI pulses during the daytime. **D)** % of *acth+* cells that are also *fosab+.* Brush handling is significantly greater than brush control (*p*=0.0047), and mesh stress is significantly greater than 6h JACUZI-SD, 6h JACUZI Day pulses, or 1h JACUZI Day pulses (*p<*0.0001). (*n* = 13, 10, 13, 5, 5, 4, 5, 6, 4,13, 11) **E)** Normalized *fosab* fluorescence intensity in *acth+* cells (each data point represents the average per animal). Brush handling is significantly greater than brush control (*p=*0.0008), and mesh stress significantly greater than 6h JACUZI-SD, 6h JACUZI Day pulses, or 1h JACUZI Day pulses (*p<*0.0001) (*n* = 10, 11, 13, 5, 5, 4, 5, 6, 4, 13, 11) **F)** % of *crh+* cells that are also *fosab+.* Brush handling is significantly greater than brush control (*p=*0.0045), and mesh stress significantly greater than 6h JACUZI-SD, 6h JACUZI Day pulses, or 1h JACUZI Day pulses (*p<*0.001) (*n* = 10, 8, 12, 5, 5, 5, 5, 6, 4,10, 8) **G)** Normalized *fosab* fluorescence intensity in *crh+* cells (each data point represents the average per animal). Mesh stress is significantly greater than 6h JACUZI-SD, 6h JACUZI Day pulses, or 1h JACUZI Day pulses (*p<*0.0001). (*n* = 10, 8, 12, 5, 5, 5, 5, 6, 4,10, 8). *(*p<0.05; **p<0.01; ***p<0.001; ****p<0.0001;* One-Way ANOVAs followed by Sidák’s multiple comparisons test)

We found that 6 hours of water pulses, applied either during the day cycle (assessing the stress of the manipulation itself apart from the stress of lost sleep) or the night cycle (assessing the combination of the stress of the manipulation and sleep loss), did not increase *fosab* in the cells of the HPI axis. Because *fosab* induction is most sensitive to changes in behavioral states, and may be diminished with such a lengthy manipulation^25^, we repeated with one hour of JACUZI pulses during the day cycle to further confirm the manipulation itself is not stressful. We still saw no significant induction of *fosab*, confirming our hypothesis that water pulses are ethologically non-threatening (Figure 4D-G). One hour of JACUZI pulses did not increase the percentage of *fosab+ pomc+* cells of the anterior pituitary (Figure 4D), nor did it increase *fosab* expression in this population (Figure 4E). However, an hour of paintbrush handling induced *fosab* in an average of 30.6±25% more *pomc+* cells compared to controls.

Furthermore, JACUZI pulses did not increase the percentage of *crh+* neurons that express *fosab*, but paintbrush handling increased the number of *fosab+ crh+* neurons by 29±22% (Figure 4F), likely because the paintbrush resembles a looming stimulus^20^. No difference was found in the intensity of *fosab* expression in *crh+* neurons with paintbrush handling (Figure 4G). Six hours of extended day did not increase *fosab* in the cells of the HPI axis. While this form of SD is minimally invasive and does not introduce the confound of stress, the introduction of a zeitgeber presents the confound of circadian manipulation for those who are interested in selectively raising homeostatic sleep pressure independent of the circadian drive for wakefulness ^19^. Together, these data suggest that JACUZI-SD does not significantly activate the HPI stress axis relative to rested controls and is also less stressful than paintbrush gentle handling.

### JACUZI-SD activates known sleep pressure pathways

To further validate our approach, we assessed whether JACUZI-SD engaged a known neural correlate of sleep pressure. Specifically, work in zebrafish and mice has connected heightened sleep pressure with induction of the neuropeptide *galanin* or activation of galanin+ neurons^13, 26, 27^. We therefore conducted HCR fluorescence *in situ* hybridization for *galanin* in the neurosecretory preoptic area (NPO) of JACUZI sleep deprived and rested controls (Figure 5A**,B**). We observed a significant increase in the fluorescence intensity of *galanin* expression induced by JACUZI-SD (Figure 5E), but not water pulses applied during the light cycle (Figure 5C**,D****,E**), suggesting that the sleep debt accumulated by JACUZI-SD replicates the neural state of sleep deprivation seen in other studies.

**Figure 5:**
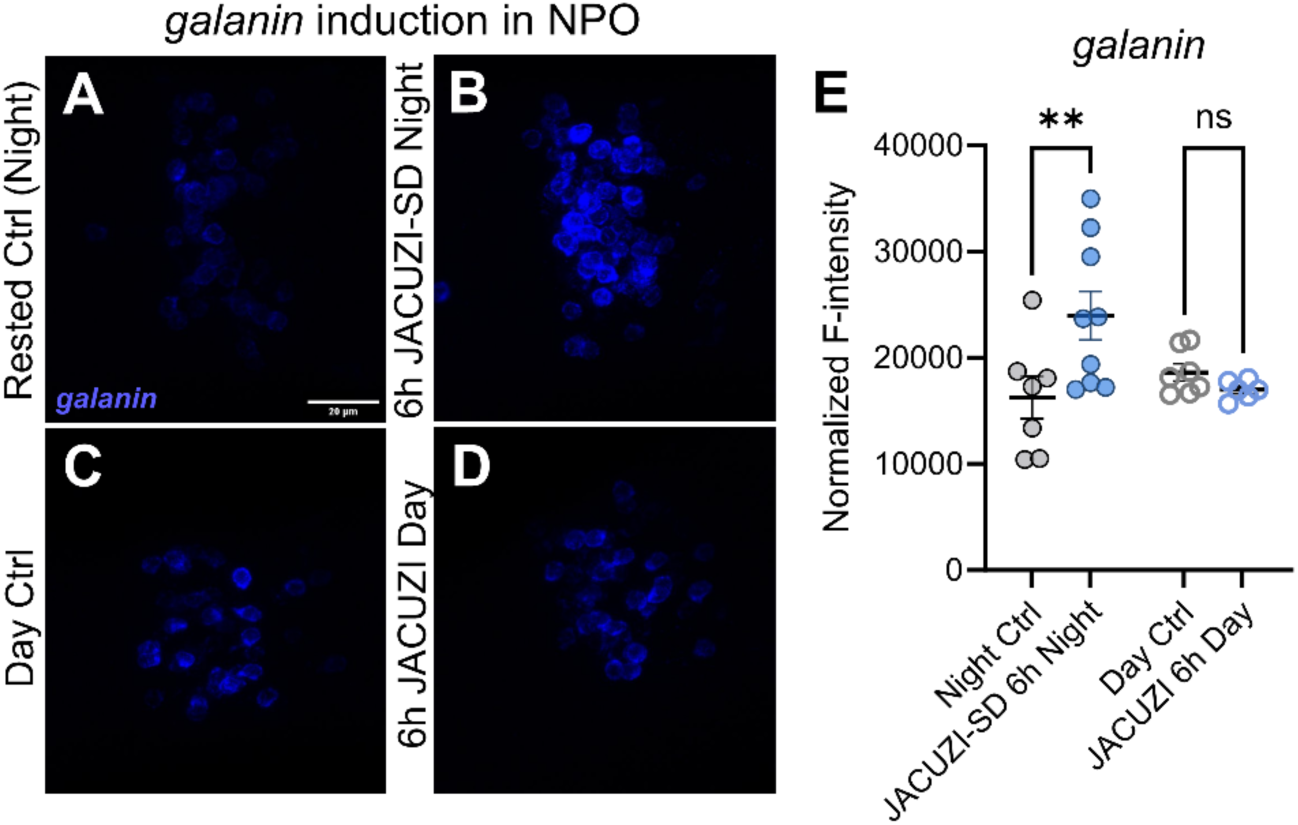
JACUZI-SD activates known sleep pressure pathways. **A-D)** Representative images of HCR fluorescent *in situs* of galanin neurons in the neurosecretory preoptic (NPO) in time-matched **A)** rested controls, **B)** clutchmate larvae after 6 hours of JACUZI-SD; **C)** daytime JACUZI controls, **D)** clutchmate larvae after 6 hours of daytime JACUZI pulses (*p=*0.0078; *n* = 7, 9, 7, 6; One-Way ANOVA followed by Sidák’s multiple comparisons test).

### JACUZI-SD as a tool to explore biological variability in homeostatic response

Homeostatically-regulated behaviors such as feeding^28^, drinking^29, 30^, sleeping^31^, and socializing^32^ are characterized by hypothalamically-driven rebound in behavior whose intensity and duration reflects the period of deprivation. For example, a greater duration of social isolation will yield a correspondingly greater rebound in social-seeking behavior^32^. Sleep deprivation by JACUZI-SD demonstrates this homeostatic response in that SD larvae exhibit a rebound sleep greater than those that rest (Figure 3). However, there is considerable variability in the homeostatic response, as some individuals exhibit a 40% greater increase in total sleep time than rested controls, while others exhibit none, or even a reduction in sleep time (Figure 6A).

**Figure 6:**
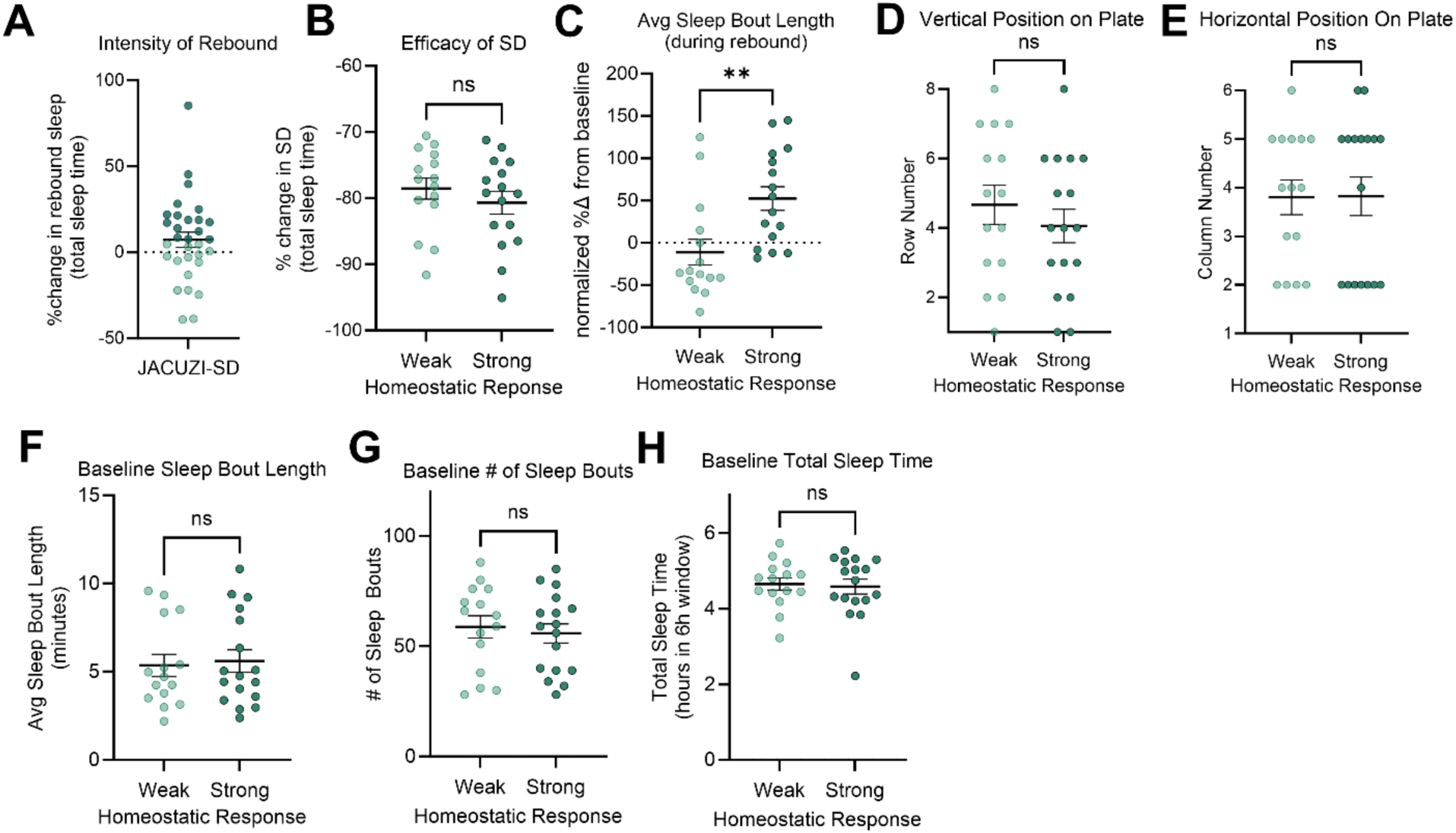
Biological variability in homeostatic rebound sleep response independent of sleep loss, technical set up, or baseline sleep microarchitecture. A) Larvae are divided into those with strong vs weak homeostatic responses – top 50% and bottom 50% of total sleep time response out of the fish that received 75% or more sleep deprivation compared to controls during two experimental replicates. B) Efficacy of sleep deprivation as measured by percent change in total sleep time from baseline sleep. C) Larvae with stronger homeostatic response in terms of total sleep time also have longer sleep bout length (*p=*0.0045). D) Row number and E) column number in JACUZI-SD plate of strong versus weak homeostatic responders. F) Baseline sleep bout length, G) baseline number of sleep bouts, and H) baseline total sleep time in strong versus weak homeostatic responders. (Unpaired t-Tests, *n* = 15, 16)

Because JACUZI-SD is unique among physical SD approaches in zebrafish in that it in enables individual sleep behavior tracking before, during, and after sleep deprivation, we could determine whether there was a relationship between the efficacy of sleep deprivation (i.e., % reduction in total sleep time compared to baseline night) and the intensity of rebound sleep (i.e., % increase in total sleep time compared to baseline night). Intriguingly, we found no relationship between the efficacy of SD and the intensity of rebound in SD animals (**Figure S5A**, R^2^ = 0.0201), but instead a positive correlation indicating individual consistency of sleep behavior throughout the night in control animals (**Figure S5B**, R^2^ = 0.2577.) With this focus on within-individual consistency in behavior, we next hypothesized that there may be natural biological variability among animals in rebound response, with some fish being “weak” and others “strong” homeostatic responders.

To test this, we separated individual data points by the top 50% and bottom 50% of rebound responses of the larvae that received 75% or greater loss of sleep. Larvae that were in the bottom 50% of rebound total sleep time we called “weak” homeostatic responders, and larvae in the top 50% of rebound total sleep time we called “strong” homeostatic responders (Figure 6A). We sought to determine whether this reflected true biological variability in rebound intensity or a methodological effect in which variability in rebound intensity corresponded to variability in efficacy of the water pulses to maintain wakefulness. Consistent with our correlation findings (**Figure S5A**), there was no difference in the efficacy of sleep deprivation between the two groups: the weak and strong homeostatic responders had lost equal amounts of sleep (Figure 6B).

To further validate our separation of the data points as biologically meaningful, we sought to determine whether other measures of rebound, like sleep bout length, were also significantly different between the groups. Indeed, strong homeostatic responders also had a 63.64% longer sleep bout length compared to weak responders (Figure 6C). We next asked whether the difference in rebound response was driven by environmental factors that differ across spatial position in the ZebraBox. However, the strong and weak responders are evenly spread across the rows and columns of the 96 well plate (Figure 6D**,E**). As the variability did not appear to be methodological or environmental, but more likely biologically driven, we wondered whether strong and weak responders exhibited inherently differing baseline sleep patterns.

Intriguingly, we observed no differences in their baseline sleep bout length, number of sleep bouts, or total sleep time (Figure 6D-F). Instead, the differences in sleep regulation are specific to the context of homeostatic rebound.

This result identifies a novel approach to understanding molecular and cellular mechanisms that regulate sleep need. The high-throughput nature of JACUZI-SD enables us to harness and explore the natural biological variability across a population. While this biological variability can be seen with other high-throughput SD approaches, such as extending daylight, JACUZI-SD uniquely offers selective manipulation of sleep pressure over circadian arousal or stressors. JACUZI-SD is thus uniquely poised to address questions of individual variability in rebound sleep among other fundamental questions surrounding the neuronal encoding of homeostatic sleep pressure.

## Discussion

The neurobiological underpinnings of homeostatic sleep pressure have remained elusive, in part because the tools for well-controlled yet high-throughput sleep deprivation are lacking. We sought a way to selectively build sleep pressure in a vertebrate model system that is both amenable to high-throughput sleep assays while also having experimentally tractable and conserved sleep circuitry. Larval zebrafish provide precisely these benefits as a model system, yielding clutches of hundreds of larvae, each with a sleep-regulatory hypothalamus that is comprised of only a few thousand neurons at four days post fertilization. To reproducibly build sleep pressure in larval zebrafish, we created JACUZI-SD: the first high-throughput, automated approach to SD in vertebrates that isolates the accumulation of homeostatic sleep pressure from common confounders such as stress or introduction of a zeitgeber.

Here we demonstrate that JACUZI-SD can induce significant levels of sleep pressure in larval zebrafish, which in turn produces substantial levels of rebound sleep. The efficacy of JACUZI-SD is further validated by induction of *galanin* expression in the NPO, an established neural correlate of heightened sleep pressure in fish and mammals ^13, 26, 27^. Furthermore, the water pulse stimuli used for JACUZI-SD are less stressful than existing SD approaches, providing a means to separate heightened sleep pressure from the confound of stress or circadian manipulation. Finally, we demonstrate the effectiveness of this platform in analyzing biological variability in homeostatic response to sleep pressure. Our findings suggest that individuals exhibit differing homeostatic responses to sleep pressure that are not driven by intensity of sleep deprivation, spatial position on the plate, nor related to baseline sleep microarchitecture.

The question of individual variability in behavioral response to a stimulus is rapidly garnering interest in the field of behavioral neuroscience^33–37^. Separating individuals into “susceptible” and “resilient” populations has led to discoveries about the neuronal underpinnings of stress^33–36^. To our knowledge, there have been few studies investigating the neural underpinnings of biological variability in homeostatic rebound sleep in vertebrate model systems. The homeostatic response to sleep deprivation is remarkably heterogeneous in the human population. Some individuals exhibit robust rebound responses to accumulated sleep debt, while others appear to need less sleep^38, 39^. Investigation of this heterogeneity has led to the discovery of sleep regulatory genes and a deeper understanding of the homeostatic regulation of sleep. We believe that JACUZI-SD, our novel high-throughput SD approach that isolates heightened sleep need from other variables, holds great potential to drive the next steps in elucidating the neural encoding of homeostatic sleep pressure.

### Limitations of the approach

While JACUZI-SD circumvents the need to separate experimental and control animals into separate tracking systems, it requires that the sleep-deprived and rested controls be on separate sides of the plate, which can introduce baseline differences in behavior if the environment is not carefully controlled. However, this can be addressed by careful control of environmental variables (see troubleshooting). Secondly, while JACUZI-SD provides sufficient SD to yield a significant rebound sleep and activate and therefore further elucidate neural underpinnings of sleep pressure, it does not provide total sleep deprivation (no sleep permitted for the entirety of the SD). There may therefore still be instances where paintbrush SD is preferable for the manual minute-by-minute confirmations of wakefulness. We expect that for many experimental questions that the SD provided by JACUZI will be sufficient and still preferable to activating stress pathways.

## Acknowledgements

We thank Frazer Matthews for animal care and members of the Mumm laboratory for consulting on zebrafish husbandry and reagents. We thank the Kanold laboratory for use of 3D printer. We thank all authors for their comments on the manuscript. This study was supported by an award from the National Institute of Mental Health (R01MH126676) to S.B., a OneNeuro Discovery Award from Johns Hopkins to S.B. and C.H., a Jane Coffins Child Postdoctoral Fellowship to L.J.E, and Wellcome Trust Investigator Award (217150/Z/19/Z) to J.R.

## Author contributions

L.J.E. and S.B. conceived the approach, and S.B. supervised all its stages. J.R. consulted at all stages of the project and initially trained L.J.E. in larval zebrafish sleep deprivation and analysis. L.J.E. designed and tested the JACUZI-SD milli-fluidics system in collaboration with H.K. and C.H. L.J.E. collected all the data, assisted by C.Z. F.K. designed behavioral data analysis pipeline, approach to excluding frames with water movement, and consulted on ZebraBox troubleshooting. L.J.E. drafted the manuscript, which all authors edited.

## Methods

### 3D Printed Mesh-Bottom 96 well plate

We used the mesh-bottomed 96 well plate CAD file reported previously ^22^ and set infill density to 50%. Depending on the individual 3D printer, infill density may need to be raised or lowered to prevent fish from escaping while allowing sufficient water flow for SD.

### Resin milli-fluidic device

The millimeter-scale fluid channels network was designed using computer aided design software (AutoCAD, Autodesk, USA). Design principle, final geometry and fabrication methodologies can be found in detail in **Supplemental Document 2**. The network was printed using a commercial stereolithography 3D printer (Form 3, Formlabs, USA) using standard resin (Clear v4, Formlabs, USA) with a 100µm layer height. Upon completion, the print was thoroughly rinsed in clean isopropanol, followed by a 5-minute sonication in a bath of fresh isopropanol, and concluded with an additional rinse using clean isopropanol for more thorough removal of uncured resin. The rinsed print was then air-dried and baked at 120°C overnight to fully cure any trace remaining uncured resin, which could otherwise be toxic to larvae or disrupt flow through the channels. Prints were gradually cooled to 25°C overnight at room temperature to limit print warping prior to use. Finally, the print was thoroughly rinsed in dH2O and air-dried overnight before use.

#### Design of the device

Two mirrored series of bifurcating millimeter-scale fluid conduits were designed to split water flow from a single pump into eight sets of six channels, each delivering uniform flow into 48 wells of a conventional 96-well plate (**Figure S1**). The mirrored arrangement of symmetric conduits was intentionally designed to enable independent operation of the two halves of the 96 wells. This ensures that stimulating larvae in either set of conduits produces synchronized waterjet performance, making responses from the left and right conduit sets indistinguishable.

Moreover, the design allows control and experimental groups to be exposed to identical environmental cues, including chemical, optical, and thermal factors, while isolating waterjet-induced arousal to one set when the other is turned off. The conduits were designed as internal channels within a 4mm tall stereolithography 3D printed block, whose lateral footprint matches a standard 96-well plate and fits beneath the 3D printed mesh 96-well plate secured in a ZebraBox chamber.

The geometry and arrangement of fluid conduit network were designed using the analogy between electric and hydraulic circuits, a common approach in constructing pressure-driven microfluidic networks ^40^. In brief, the flow rate through each conduit segment under a set pumping pressure was estimated based on the hydraulic resistances, as determined by the segment geometry and fluid viscosity. This approach leverages the analogy of flow rate as electrical current and applied pressure as potential difference in electrical circuitry. The fluid conduit network consists of rectangular channels arranged in parallel and series to deliver the set amount of flow rate, where hydraulic resistance,^40^ Rh, is calculated as:

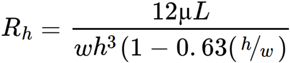

Here, L, w, and h represent the length, width, and height of each fluid conduit segment, respectively, and µ represents the dynamic viscosity of water. To connect to the water pump, standard ¼” female National Pipe Taper (NPT) fittings were 3D printed onto the device to connect to commercially purchased adaptors.

### Assembly of JACUZI device

Epoxy was applied to the mesh bottom of the 96 well plate using a pipet tip, tracing the top, bottom, and internal side of each half of the plate. The epoxy thus formed a “C” shape, creating a barrier between the left and right sides of the plate that extends to trace the top and bottom edges while leaving the far right and far left edges open (Figure 1G). This prevents the control fish from experiencing water level disturbances during SD.

### Zebrafish

WT AB embryos were staged and sorted at 4-7 hours post fertilization and kept in a reverse light-dark cycle incubator at 28C +/-1C. Lights turn on at 7:30pm and off at 9:30am. The morning of 4dpf, before the lights went off, fish were screened for swim bladders, and those without fully inflated swim bladders were removed.

### Arduino Controlled Submersible Water Pump

We programmed the submersible water pump (Model # 3-01-1249, HiLetgo) to an Arduino, allowing precise and randomized control of pulse frequency and duration. The pump is rated at 0.16 gallons/minute (167 mL/s) by the manufacturer, and we measured a comparable bulk flow rate of 100.0 ± 2.3 mL/s during short pulse durations (<1s) ranging from 100 to 250ms (**Figure S6**). Random number generators embedded in the Arduino code were used to vary pulse duration (100-250 ms) and inter-pulse interval (1-25 s), helping to reduce habituation to repetitive stimuli. Over the six hours of stimulation, the pulse duration increased and the inter-pulse interval decreased each hour. In this way, the stimulation was gradually intensified to combat the increasing sleep pressure, such that the intensity was not excessive and stressful for the start of SD but was still sufficient to maintain wakefulness in the sixth hour. Of note, the specific ranges of pulse durations were adjusted per experiment to reduce the risk of overflow or cross-well spillage. The Arduino Code with specifications is available in **Supplemental Document 1**. Wiring diagram and beginner Arduino tutorial can be found here: https://arduinogetstarted.com/tutorials/arduino-controls-pump.

### Setting up the ZebraBox for JACUZI-SD

Two lengths of 3/8” ID tubing were threaded through the back panel of the ZebraBox (ViewPoint LifeSciences). 3/8” threaded stainless steel barb adapters were screwed into the milli-fluidic device ports. Parafilm was used to seal the connection between stainless steel adapters and milli-fluidic ports to minimize loss of water flow. The 3/8” ID tubes that were threaded into the ZebraBox were then fitted over each barb, and additional parafilm was used to seal the connections between barbs and tubing. The plate was fitted into the ZebraBox plate holder and leveled.

A ∼1” length of ¼” tubing was used as a drainpipe in one of the outlets of the ZebraBox water bath. The height of the drainpipe should be equal or just below the height of well plate to allow water to drain before the well plate overflows, but not earlier. Parafilm was used around the tubing base to promote a water-tight seal so water only drains when excessively high. The other outlet of the ZebraBox water bath was plugged with parafilm and similar tubing that is excessively tall so that it never allows water to drain.

The other end of the tubing was connected to a 2” length of 5/16” diameter tubing which functioned as an adaptor to attach to the outlet port of the submersible water pump. Parafilm was used to keep the connection between the tubing and the pump watertight. The control side of the plate was backfilled first: the water pump, attached to the tube that feeds the control/rested side of the plate, was submerged in a 2L beaker of E3. The water pump was supplied power until the control channels and plate were filled. Then, the tubing fitted to the SD side of the plate was attached to the water pump and similarly backfilled.

Water levels were adjusted so the meniscus was 1-2mm beneath the rim of each well. Because both SD and control wells are open (i.e., not made water-tight with epoxy) to the water bath on both lateral sides of the plate (**Fig 1G**), the water levels equilibrate throughout the entire plate without a continuous recirculation device via the principle of communicating vessels.

Water pressure from the external reservoir maintains passive water influx via the two tubes – one attached to the SD side, one attached to the control side. If water levels are too low because of evaporation, water is passively drawn through the tubes. If water levels are too high, such as when the pump is supplied power during the pulses, excess water above the height of the wells drains via drainpipe. A photograph of the setup is supplied in **Figure S4.** Passive water level maintenance via the principle of communicating vessels avoids disruptive or frequent water level top-ups, and avoids the need for constant water recirculation. If excessive evaporation occurs, water levels can be raised by simply pouring water into the external reservoir until the water levels equalize with the 96 well plate to the desired level.

A wide-mouth plastic transfer pipet was used to transfer individual larvae into the 96 wells, alternating left (SD) and right (control) sides, so as not to bias the behavioral groups. In the bottom left well, a small piece of fabric was placed instead of a larva. This served as a “sentinel fish” to detect when the water pulses were actively causing movement (**Figure S2, Supplemental Video 1**).

A HOBO Pendant Light/Temperature Logger (Onset) was placed in the water bath to ensure temperatures remained 26-29C throughout the experiment. The ZebraBox was closed and the front was wrapped in light-blocking fabric. Additional light-blocking fabric was secured around the tubing where it exits the back panel of the ZebraBox. An acoustic foam cover was placed over the entire ZebraBox to further minimize light and sound pollution.

### Experimental Timeline

A standard sleep experiment was set up in the ZebraLab software using a 14hr:10hr reverse light-dark cycle, lights off at 9:30AM and lights on at 7:30PM. Night 1 (4dpf) and day 1 were considered habitation periods; night 2 (5dpf) and day 2 were used as baseline datasets; and the sleep deprivation pulses occurred for the first 6 hours of night 3 (6dpf), while the last 4 hours of night 3 were used as recovery or rebound sleep. The experiment concluded at 9:30AM the morning after day 3.

After the experiment, the larvae were euthanized in ice water. The plate was rinsed with dH2O and submerged in 5% bleach for 2-4 hours. The plate was thoroughly washed five times with dH2O by flushing both sets of channels and fully submerging, and rocking, the plate in dH2O. The plate was kept submerged in dH2O overnight and then given a final rinse in dH2O before being left to dry completely for 1-2 days before beginning another experiment.

### Sleep Deprivation by Water Pulses

The Arduino script, which determines the pulse duration and frequency for each hour of sleep deprivation, is uploaded to the Arduino. The pump is supplied power by plugging the 12V adaptor into an outlet. Pulses are watched carefully for a few minutes to ensure they are causing water disturbances in each well without flooding water over the wells.

### Collection, Normalization, and Statistical Analysis of Sleep-Wake Behavior

After exporting the ZebraLab results file and exporting the ZebraLab raw data files, the FramebyFrame R package (https://github.com/francoiskroll/FramebyFrame) ^22^ was used for the analysis. Pulse frames, detected using the “sentinel fish”, were replaced by Δpx = 0 (**Figure S2**). The function first recorded the sentinel fish’s “active bouts” during the JACUZI-SD period, defined by any continuous series of positive Δ pixel values lasting at least 2 frames and reaching Δpx = 2. Each recorded pulse was extended by 3 frames (0.12 sec) in the past and 2 frames (0.08 sec) after the last positive value to include potential water movement that did not move the sentinel fish. In the data of the fish undergoing JACUZI-SD, the corresponding frames were replaced by Δpx = 0. FramebyFrame then calculated total sleep time, sleep bout length, sleep bout number, percentage time active, and number of active bouts for a given indicated time window.

Most data are displayed as normalized percent change from baseline as a way of controlling for both individual variability and developmental changes in sleep behavior. We first calculated percent change from each animal’s individual baseline data during the equivalent time window 24 hours prior to control for individual variability in sleep behavior. The average percent change across control animals was then subtracted from all values for a given experiment to normalize/correct for developmental changes that occur over the 24 hour period. For example, total sleep time decreases on average 20% in control animals from the 5 dpf baseline night to 6 dpf JACUZI night, but 61-84% in JACUZI-SD animals, so normalized data are presented as 41-64% reduction in sleep time beyond what is observed in rested controls. An unpaired t-test was used to detect significant differences between these values calculated from SD and rested control groups. If greater than two experimental groups were compared, a One-Way ANOVA was used to evaluate overall significance across an experiment followed by multiple comparisons between groups if so. Data points three standard deviations away from the mean were excluded as statistical outliers.

### Troubleshooting JACUZI-SD

Before introducing a new material, especially plastic, it is best to assess whether it might be toxic to larvae by incubating the material in E3 with larvae for several days. Clean plates with dilute (5%) bleach and rinse thoroughly in dH2O, as larvae are sensitive to trace amounts of bleach.

The water height in the beaker will match water height in the ZebraBox bath. If you need water levels to rise in the 96 well plate, you can add water to the beaker or raise the beaker on pieces of cardboard. Keep this in mind when optimizing control of water levels throughout the experiment, as the rate of evaporation may be different in the beaker compared to inside the ZebraBox. Our system seemed to be optimally balanced by loosely covering the beaker in plastic Saran wrap.

Maintaining even temperature and humidity in the room will ensure constant water levels, and water level changes, from one experiment to another. Pulse duration and frequency may need to be optimized to the nuances of a particular set up. For example, if the water is draining slowly, our pulse script may cause the plate to flood. Allowing water to freely drain by clearing excess epoxy, or adjusting the size of the mesh, may help. You may also try completing a full rectangular barrier around the 48 wells on the left and the right. This should maximize water flow going to the fish, though it may slow drainage. Generally, aim to choose the shortest pulse duration or lower pulse frequency to maintain wakefulness and induce rebound sleep while avoiding floods. If tubing interferes with camera view of wells, additional tape can be used to hold back the tubing. If the plate fails to sit securely, securing the tubing within the box with additional tape can also help hold the plate in place.

### HPI-axis activation

This experiment was conducted in the light phase to measure the stress contributions of manipulation alone, independent of the stress of sleep deprivation. To induce mesh stress: a mesh cell strainer containing larvae was removed and submerged from a dish of E3 every 30-90 seconds for 30 minutes (adapted from previously published work^23, 24^). For paintbrush gentle handling: larvae were plated in a 96 well plate and allowed to habituate for four hours before manipulation. The paintbrush was inserted into each well approximately once per minute, as described ^11^, for one hour. As with gentle handling used to sleep deprive larvae, care was taken not to touch the larva unless they failed to move, in which case the tail was gently approached and tapped until movement was observed ^11^. Paintbrush controls were similarly plated but placed aside on the same benchtop and left undisturbed for the hour. JACUZI fish were exposed to one hour of JACUZI water pulses, while JACUZI controls were undisturbed in the rested control side of the plate in the ZebraBox. At the end of each manipulation, larvae were euthanized by tricaine overdose (>0.3mg/mL) and immediately fixed in 4% paraformaldehyde before proceeding with HCR as described below.

### Galanin HCR

Immediately following the 6 hours of water pulses had completed during a standard JACUZI-SD experiment, JACUZI-SD larvae and rested controls were euthanized by tricaine overdose and immediately fixed in 4% paraformaldehyde. HCR staining was completed according to Molecular Instruments’ *HCR v3.0 protocol for whole-mount zebrafish embryos and larvae* with the following modifications: brains were dissected after an initial overnight (15-18h) fixation followed by PBS washes, and no methanol dehydration steps were performed (embryos went straight from dissection in PBS to PBST (0.1% Tween-20 in PBS) washes). Larvae were mounted whole in 1.5% low-melt agarose in an 8 well chamber slide for confocal imaging on a Zeiss LSM 700.

### Analysis of Confocal Images

Fluorescence intensity was measured by blinded scorer in Fiji. The average fluorescence intensity per animal was calculated as the average fluorescence intensity of cell bodies and then normalized to background fluorescence of the same image. A cell body was considered *fosab+* if, at a minimum, there were notably more mRNA fluorescence dots than background, typically >3-5 dots per cell body. One-Way ANOVA was used to evaluate overall significance across an experiment followed by multiple comparisons between groups if so. Data points three standard deviations away from the mean were excluded as statistical outliers.

---

**Supplemental Figure 1:**
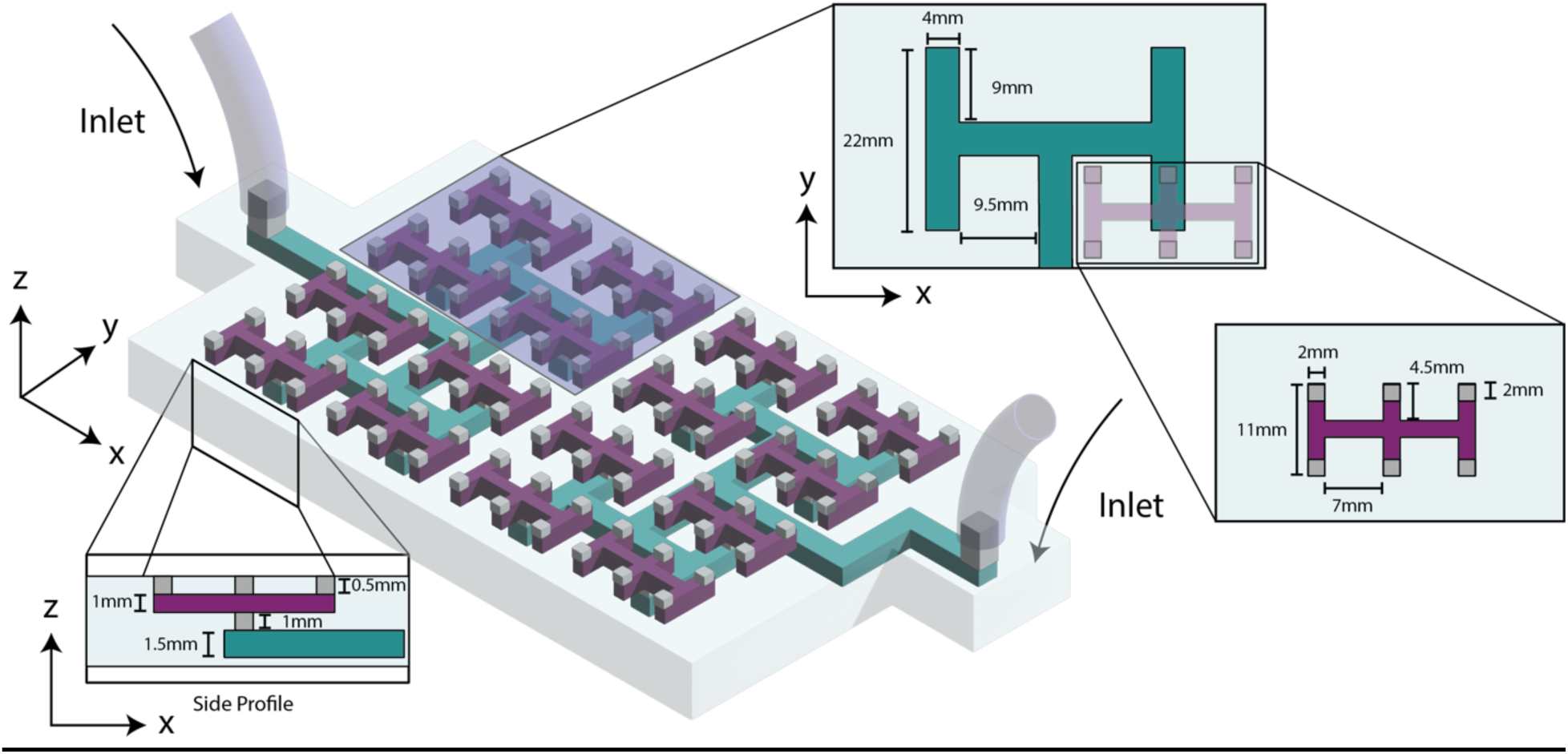
Schematic of the mirrored, 3D printed milli-fluidic channel network designed to inject uniform water-jets into 48 wells. Water is pumped into the first-level channel network, which is divided into 8 segments (teal). Each segment is connected to the second-level channel network, which splits into 6 daughter channels (purple). The water then flows out of 2mm square outlets centered at the floor of each well where larvae reside.

**Supplemental Figure 2:**
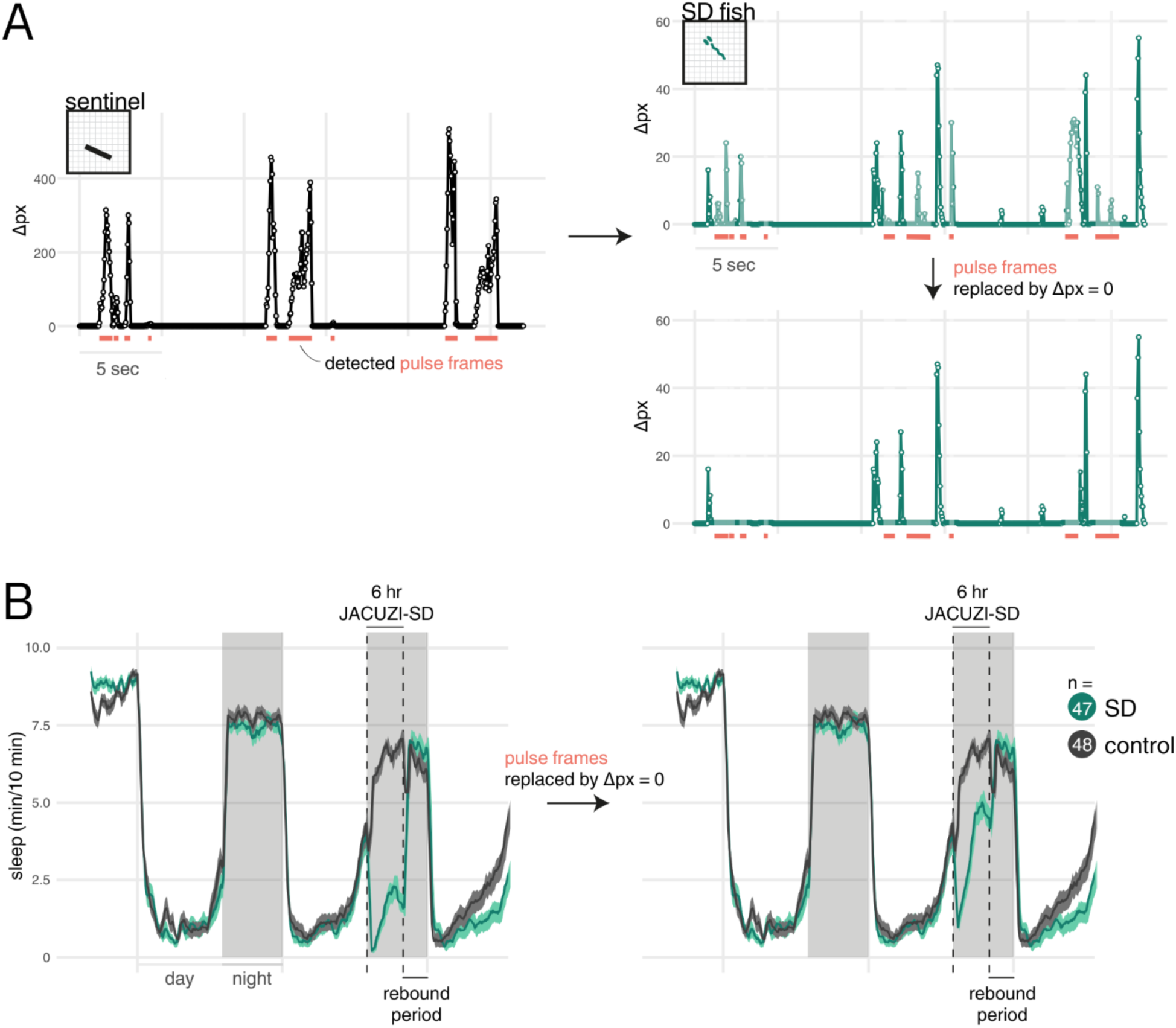
Pulse frame exclusion to generate a conservative estimate of the minimum sleep deprivation achieved. **(A)** (left) A black piece of fabric serves as a “sentinel fish” to detect frames during which the pulses moved the water. (right) For the fish undergoing JACUZI-SD, these frames are excluded by replacing them by Δpx = 0, making the assumption that any movement recorded was not genuine wakeful activity. **(B)** Sleep (minutes per 10-minute epoch) during 67 hr on a 14 hr:10 hr light:dark cycle, before (left) and after (right) pulse frame exclusion for the fish undergoing JACUZI-SD. See also Supplemental Video 1.

**Supplemental Figure 3:**
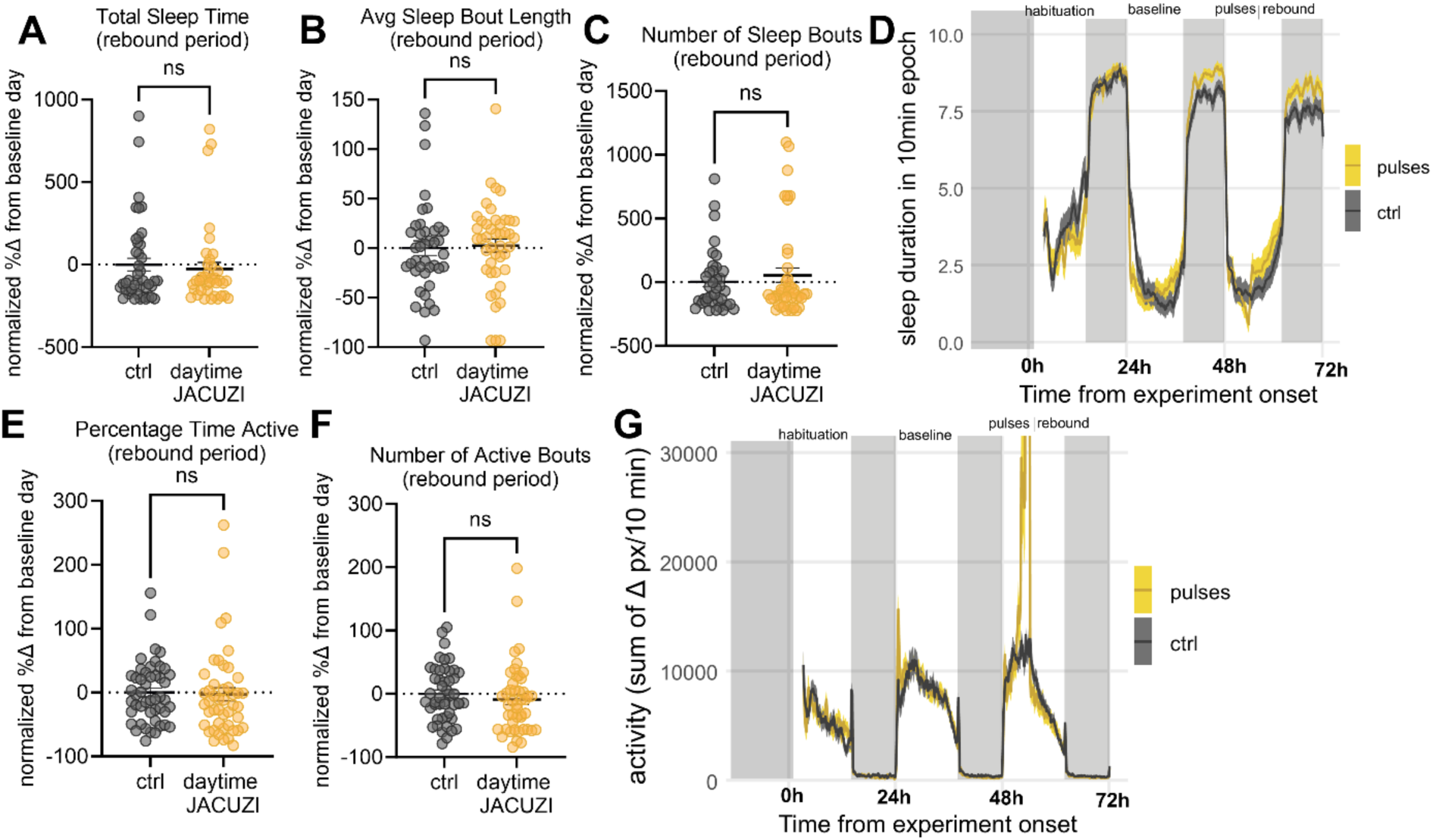
JACUZI-SD water pulses do not induce rebound when applied during the light cycle. Normalized percent change in A) total sleep time (*p=*0.6359; *n* = 43, 39), B) sleep bout length (*p=*0.7950; *n* = 40, 44), C) number of sleep bouts (*p=*0.4630; *n* = 37, 43) during the 7h rebound period compared to the 7h baseline period 24h prior. Average sleep trace shown in D. Normalized percent change in E) percent time active (*p=*0.8133; *n* = 48, 46), and F) number of active bouts (*p=*0.3782, *n* = 48, 46) during the 7h rebound period compared to the 7h baseline period 24h prior. Average activity trace shown in G. *(*p<0.05; **p<0.01; ***p<0.001; ****p<0.0001;* Unpaired t-Tests)

**Supplemental Figure 4:**
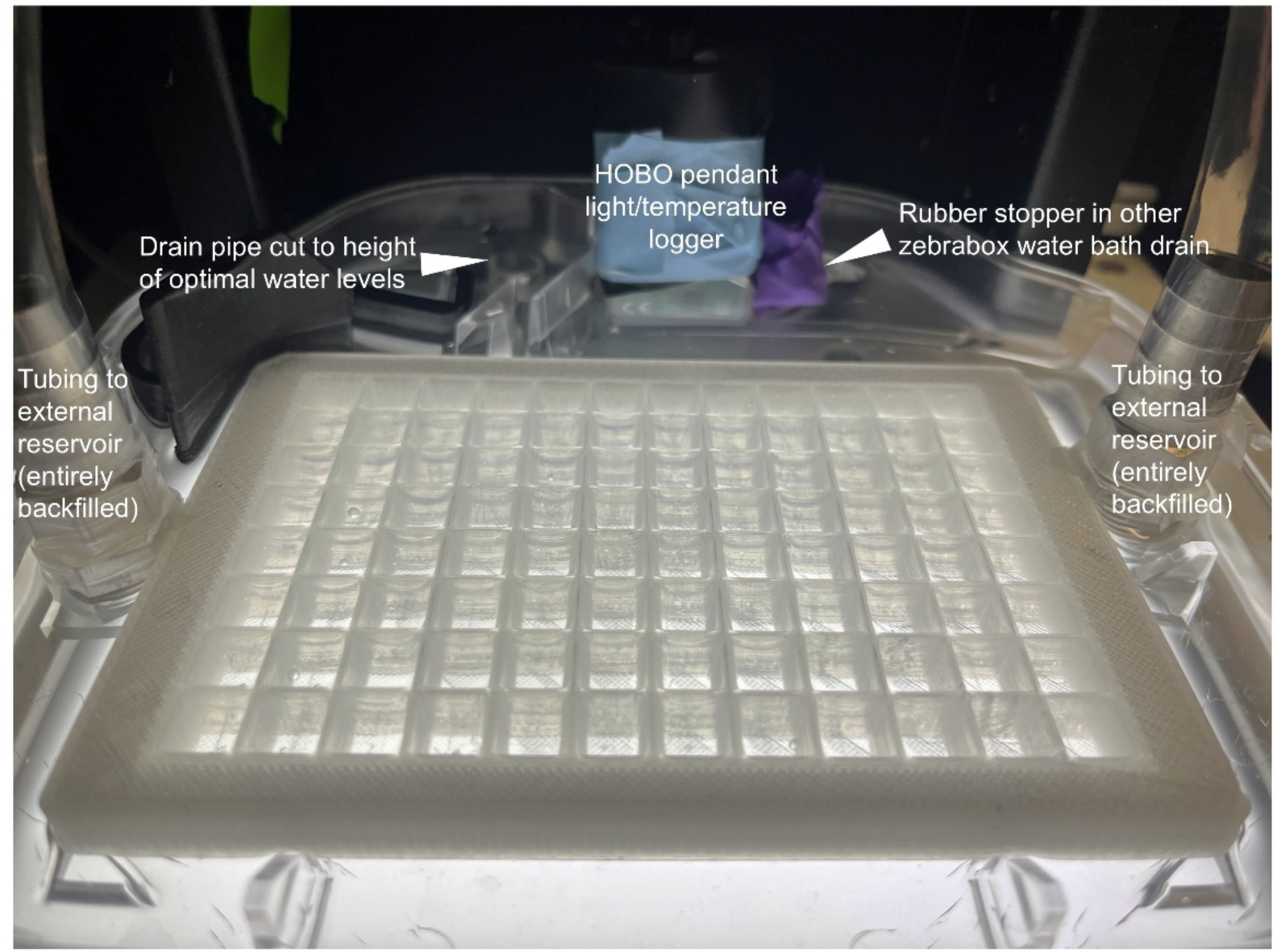
Photograph of JACUZI-SD set up inside the Zebrabox to demonstrate how water levels are maintained.

**Supplemental Figure 5:**
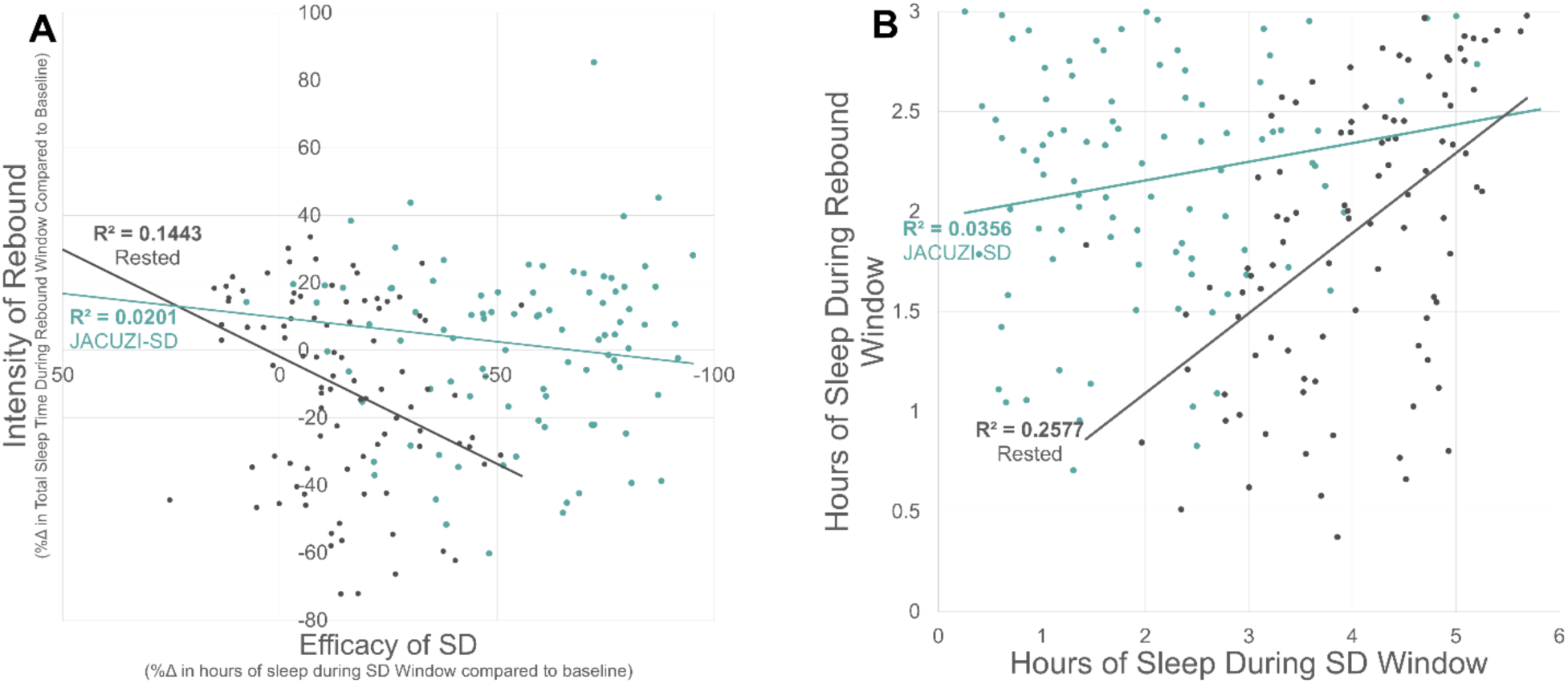
Within-individual consistency in sleep behavior is a stronger predictor of future sleep behavior than efficacy of SD. **A)** There is no correlation between efficacy of SD and intensity of rebound (R^2^=0.0201). **B)** Control fish exhibit a positive correlation between total sleep time at beginning and end of 6dpf night, suggesting within-individual consistency in sleep pattern is strong (R^2^=0.2577). Fish that sleep a lot at the start of the night will sleep a lot at the end of the night; fish that don’t sleep very much at the beginning of the night don’t sleep very much at the end of the night. This relationship is greatly weakened when we sleep deprive (R^2^=0.0356), but not to the point of a negative correlation: those that sleep less during SD don’t necessarily exhibit more rebound.

**Supplemental Figure 6:**
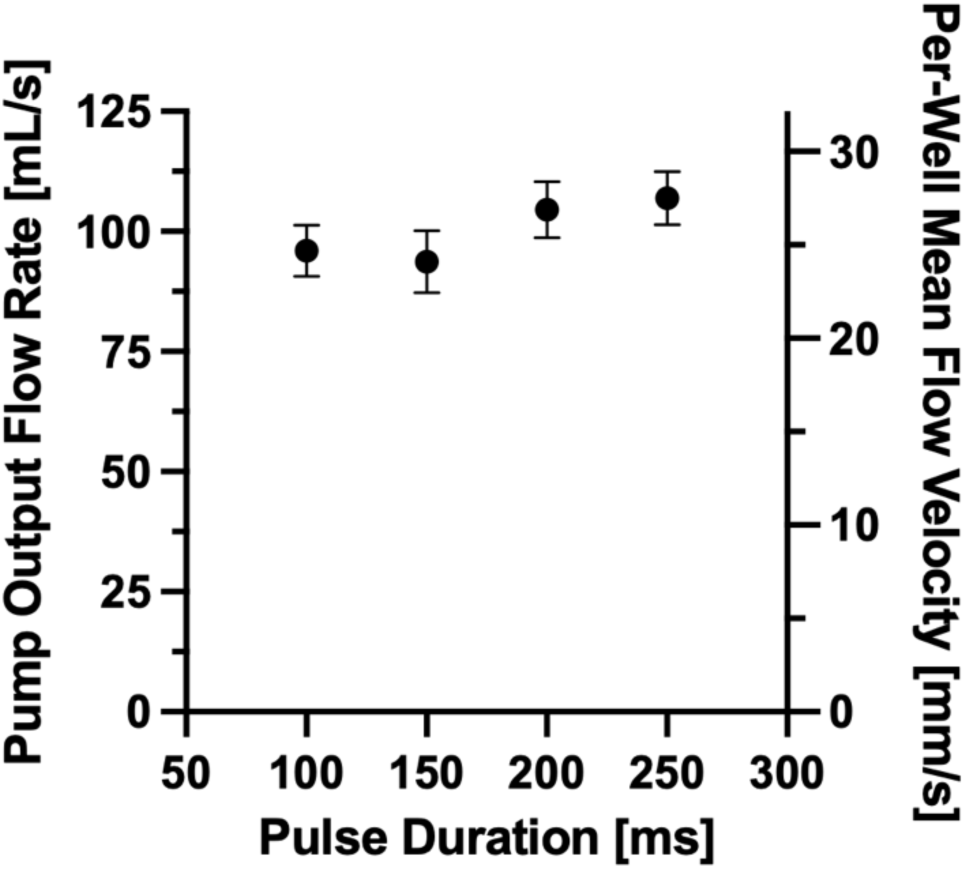
Pump output flow rate measurements from the Arduino controlled water pump across pulse durations of 100-250 ms. Flow rates were consistent across conditions (p>0.2) and comparable to the reference rate measured with a 1 second pulse (102.8 ± 0.4 mL/s). Average per-well flow velocity was estimated assuming uniform distribution across 48 outlets feeding into 9mm square wells. Values represent mean ± SEM, averaged over 3 independent replicates.

**Supplemental Video 1**: Three fish are depicted during a single water pulse. There are two types of movements: volitional movement of the fish before and after the water movements (during Pulse OFF), and involuntary swirling of the fish as the pulse generates water movements (during Pulse ON). Without pulse frame exclusion (**Figure S2**), the frames in which the fish are involuntarily moved by the water are counted as wakeful, thereby including some potentially false positive wakefulness. With pulse frame exclusion, the frames during the water movements are not counted as wakeful movements, but exclusively the unambiguously volitional movements outside this time window. These time windows are calculated by including a sentinel fish in each experiment, a black piece of fabric roughly the same size as a larva, which generates movements during the water movement but not outside the water movement (See **Figure S2** and Methods for details on calculations).

**Supplemental Document 1:** Arduino Code for JACUZI-SD pulses.

**Supplemental Document 2:** CAD file for stereolithography millifluidic device.

