## Supplemental Document 2 for "JACUZI-SD: An automated, high-throughput, minimally stressful approach to sleep depriving larval zebrafish": Elias et al_Supplemental Document 2.ino

```
/*
 * Created by ArduinoGetStarted.com
 *
 * This example code is in the public domain
 *
 * Tutorial page: https://arduinogetstarted.com/tutorials/arduino-controls-pump
 */

// constants won't change
const int RELAY_PIN = A5;  // the Arduino pin, which connects to the IN pin of relay
long randOn;
long randPause;
long i = 1;
// the setup function runs once when you press reset or power the board
void setup() {
  // initialize digital pin A5 as an output.
  pinMode(RELAY_PIN, OUTPUT);
  randomSeed(analogRead(0)); //generate different seed numbers each time the sketch runs, only works if analog input pin 0 is unconnected
  }

// the loop function runs over and over again forever
// this is the SD function
void loop() {   
  //depending on how long the SD has been going, determine the pulse duration and delay before pump goes on again
  //if it's been going for less than one hour (3600000)
  if(i>0 && i<3600000){
  randOn = random(100, 120); //generate a random number - duration for ON between 100 and 120 ms
  randPause = random(12000, 25000); //generate a random number - duration for OFF between 12-25 seconds
  }

  //if it's been going for more than one hours but less than two hours (7200000)
  else if (i>3600000 && i<7200000){
  randOn = random(110, 150); //generate a random number - duration for ON between 110 and 150 ms
  randPause = random(10000, 17000); //generate a random number - duration for OFF between 10-17 seconds
  }

 //if it's been going for more than two hours but less than three hours (10800000)
  else if (i>7200000 && i<10800000){
  randOn = random(120, 170); //generate a random number - duration for ON between 120 and 170 ms
  randPause = random(7000, 12000); //generate a random number - duration for OFF between 7-12 seconds
  }

  //if it's been going for more than three hours but less than four hours (14400000)
  else if (i>10800000 && i<14400000){
  randOn = random(120, 190); //generate a random number - duration for ON between 120 and 190 ms
  randPause = random(3000, 10000); //generate a random number - duration for OFF between 3-10 seconds
  }

//if it's been going for more than four hours but less than five hours (18000000)
  else if(i>14400000 && i<18000000){
  randOn = random(130, 220); //generate a random number - duration for ON between 130 and 220 ms
  randPause = random(2000, 9000); //generate a random number - duration for OFF between 2-9 seconds
  }

  //if it's been going for more than five hours but less than six hours (21600000)
  else if(i>18000000 && i<21600000){
  randOn = random(140, 250); //generate a random number - duration for ON between 140 and 250 ms
  randPause = random(1000, 7000); //generate a random number - duration for OFF between 1-7 seconds
  }

  //if it's been going for more than six hours - delay 24 hours (stop SD)
  else if(i>21600000){
    digitalWrite(RELAY_PIN, LOW);  // turn off pump
    delay(86400000);
  }

  digitalWrite(RELAY_PIN, HIGH); // turn on pump
  delay(randOn); // keep on for a random duration from 100 to 150 milliseconds
  digitalWrite(RELAY_PIN, LOW);  // turn off pump 
  delay(randPause); // keep OFF for a random duration determined by above conditionals
  //delay(6000);
  i = i + randOn + randPause; // i will keep track of the total number of milliseconds elapsed since beginning this SD loop. Keep storing the delays.
  }
```
